# Dissociable roles of cortical excitation-inhibition balance during patch-leaving versus value-guided decisions

**DOI:** 10.1101/2019.12.15.877019

**Authors:** Luca F. Kaiser, Theo O.J. Gruendler, Oliver Speck, Lennart Luettgau, Gerhard Jocham

## Abstract

In a dynamic world, it is essential to decide when to leave an exploited resource. Such patch-leaving decisions involve balancing the cost of moving against the gain expected from the alternative patch. This is in contrast with value-guided decisions that typically involve maximizing reward by selecting the current best option. Patterns of neuronal activity pertaining to patch-leaving decisions have been reported in the dorsal anterior cingulate cortex (dACC), whereas competition via mutual inhibition in the ventromedial prefrontal cortex (vmPFC) is thought to underlie value-guided choice. Here, we show that the balance between cortical excitation and inhibition (E/I balance), measured by the ratio of GABA and glutamate concentrations, plays a dissociable role for the two kinds of decisions. Patch-leaving decision behaviour was related to E/I balance in dACC. In contrast, value-guided decision making was related to E/I balance in vmPFC. These results support previous mechanistic accounts of value-guided choice and provide novel evidence for a role of dACC E/I balance in patch-leaving decisions.

## Introduction

In an ever-changing world with non-uniformly distributed goods, organisms have to decide whether they want to accept the resources provided by their current environment or switch to an alternative course of action. These patch-leaving decisions require balancing potential benefits in alternative environments against costs associated with abandoning the current patch (or current course of action). Patch-leaving decisions can be contrasted with value-guided choices, where agents often need to integrate multiple attributes to select the option currently most valuable. Consider for example a researcher working at a university in a small town who considers moving to Munich (famous in Germany for high cost of living). Initially, there is no benefit in leaving: Moving costs money and the cost for living is higher in Munich. However, better career prospects and a higher salary might, in the long run, overcompensate the financial and social costs incurred. This constitutes a patch-leaving decision. In Munich, our researcher may face a decision on where to live - and attributes like the quality of different apartments, the rent, and the distance from the office may contribute to how valuable each flat is judged. Based on these attributes, our researcher would simply select the one they judge more valuable altogether, thus maximizing immediate reward. This kind of decision is commonly referred to as a value-based decision.

Studies in animals and humans suggest a role of the dorsal anterior cingulate cortex (dACC) in patch-leaving decisions^1–4^, as well as signaling potential costs entailed by behavioural adjustments^2, 3^. Activity in the dACC has been reported to reflect diminished rewards within the current environment^2, 5, 6^ as well as the average value of potential alternatives^3^ suggesting an important role in guiding behavioural adjustments^7^. In contrast, value-guided decision making has been linked to the ventromedial prefrontal cortex (vmPFC)^8–16^. Activity in this region covaries with the values of the available options, positively with the value of the chosen option, and negatively with the value of the unchosen option^8–10, 14^. Both theoretical and experimental results strongly suggest that a mechanism based on competition via mutual inhibition in vmPFC supports value-guided choice^15, 17^. This competition is driven by the balance between GABAergic inhibition and recurrent glutamatergic excitation. Concentrations of the major excitatory and inhibitory neurotransmitters, glutamate and GABA, have been shown to be related to both choice performance and a vmPFC value comparison signal in a manner that is consistent with biophysical models^9, 17^.

Based on earlier reports^4, 18^, we hypothesized that patch-leaving behaviour is guided by the balance between cortical excitation and inhibition (E/I balance) in dACC. In contrast, we expected that value-guided decision making is governed by E/I balance in vmPFC. Healthy human participants performed a novel decision making task (Figure 1A) combining patch-leaving and value-based decision-making. We measured GABA and glutamate concentrations using magnetic resonance spectroscopy (MRS) at 7T in five cortical areas of interest: vmPFC, dACC, dorsolateral prefrontal cortex (dlPFC), and bilateral primary motor cortex. We report contributions of E/I balance that were dissociable as a function of decision type and cortical area. Patch leaving behaviour was related to E/I balance in dACC, but not in any of the other regions investigated. In contrast, value-guided decision making was related to E/I balance in vmPFC, but not in any of the other cortical areas.

**Figure 1.**
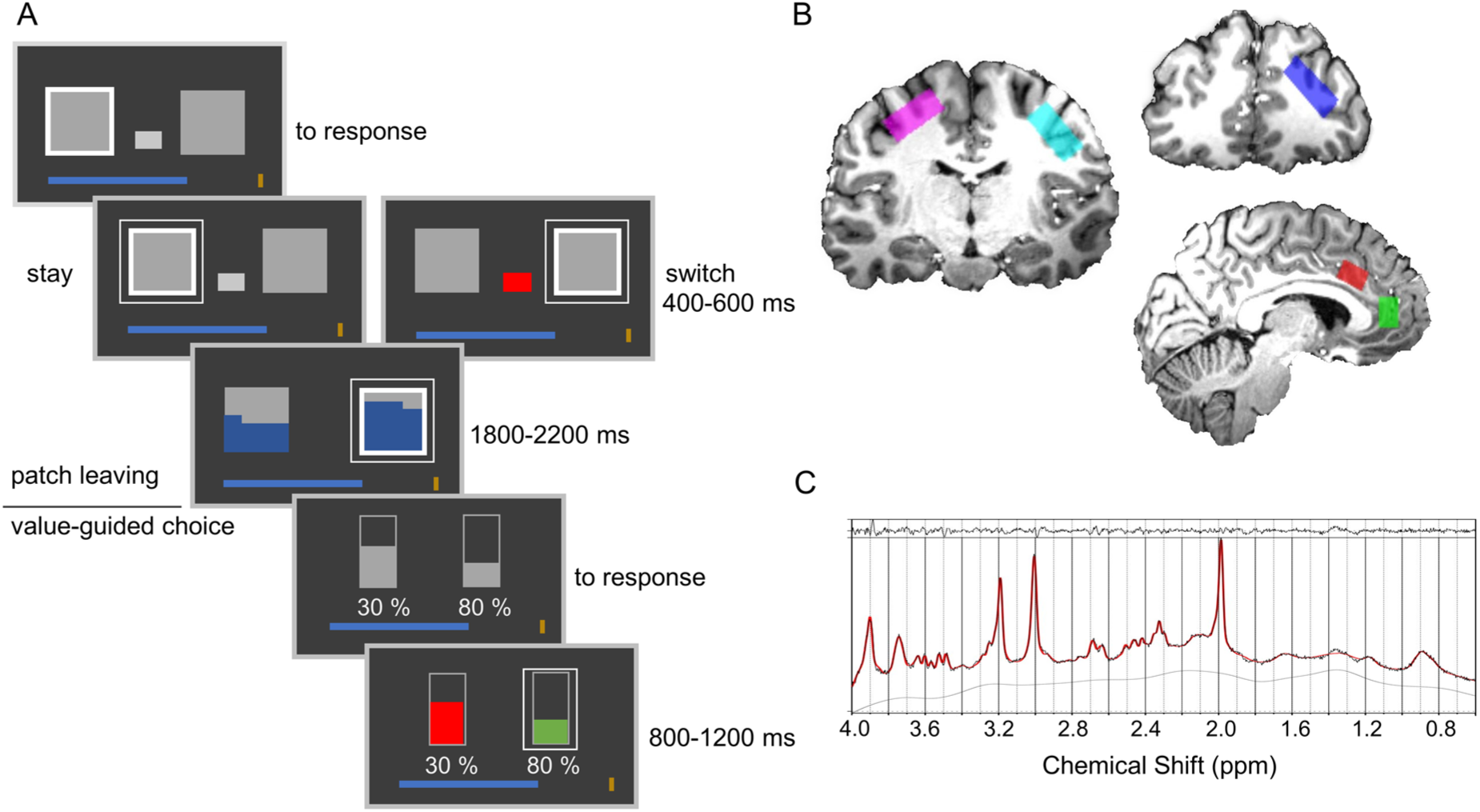
Behavioural task and MRS recordings. A) Schematic of the task structure. Participants made a patch decision in the first stage of each trial. A white outline indicated the location of the participant’s current patch. If they chose to switch, they had to pay a cost indicated by the size of a grey rectangle. After choice, the values of both patches were revealed, indicated by the blue filling. In the value-guided choice phase, the chosen patch value was randomly divided up between the two different options and assigned with random reward probabilities. Participants selected an option by pressing a button, which was followed by feedback on both options. If they obtained a reward, the blue progress bar at the bottom of the screen grew in proportion to that reward. B) Example MRS Voxel placement for one participant. Spectroscopy voxels were placed in right dlPFC (blue), bilateral motor cortices (pink and cyan), dACC (red) and vmPFC (green). C) Spectrum obtained from dACC of one exemplary participant.

## Results

Participants performed 320 trials of a novel behavioural task combining patch leaving and value-guided choice. Each trial of the behavioural task consisted of a patch leaving decision followed by a value-guided choice (Figure 1A). Importantly, the task was designed such that the value-guided choice was explicitly temporally separated from the choice to leave or stay in the current patch. At the first stage, participants indicated by button press whether they wanted to stay in their current patch or leave for the alternative patch. Leaving was associated with a cost (randomly drawn from the set {5, 10, 15, 20}) which was subtracted from the participant’s current total earnings. Over trials, the reward available in the current patch stochastically depleted according to a decaying Gaussian Random Walk, whereas the reward in the alternative patch replenished. The cost level was displayed to participants and remained constant until a decision to leave the patch was made, at which time a new cost level was randomly selected. Thus, participants needed to monitor, over trials, the relative advantage of leaving for the alternative patch and to compare this against the cost for leaving. No money could be won at this stage. Following the patch decision stage, participants entered the value-guided choice. Here, the reward available in the patch chosen by the participant was randomly divided and allocated to two choice options. Additionally, a probability with which this reward could be obtained was randomly assigned to each of these two options. This design feature ensured that the value-guided choice was temporally decorrelated from the choice to leave or stay in the current patch. While being in a rich patch will, on average, lead to better choice options at the value-guided choice stage, the exact options to choose from are not known to participants when they make their patch choice. After choice, participants received a feedback on whether their choice had been rewarded. This was followed by the next trial. In a separate session, 24 to 48 hours after volunteers completed the behavioural task, we obtained estimates of GABA and glutamate concentrations in five cortical areas of interest (Figure 1B) using single-voxel magnetic resonance spectroscopy (MRS) at 7T (see figure 1C for an example spectrum from one participant). We recorded from the dACC, the vmPFC, the right dorsolateral prefrontal cortex (dlPFC) and the bilateral primary motor cortices (M1). Note that the vmPFC voxel is located in a rather dorsal position, covering parts of pregenual ACC. This location, which is also in line with previous work^9^, was chosen based on methodological considerations since obtaining MRS measurements in more ventral positions is difficult due to field inhomogeneities. However, please also note that value signals in vmPFC, while not centred on this location, often extend to cover this region across a large swath of the ventral to dorsal extent of the mesial prefrontal cortex^9^. In addition to vmPFC and dACC, we selected the dlPFC because of its importance for working memory-related processes^19, 20^. Since patch leaving requires carrying a representation of patch leaving value across trials, we reasoned that dlPFC E/I balance might play a role in patch-leaving, but not value-guided choice behaviour. The motor cortex was selected as a control region, where we expected a relationship with motor, but not task-specific parameters, neither value- nor patch-leaving-related.

### Patch-leaving behaviour

Participants took costs and patch value differences into account in guiding their patch-leaving choices. On average, participants left their current patch on 20.55 ± 1.40 (mean ± SEM) out of 320 trials. We found that participants stayed longer in their current patch when they had to pay higher switch costs. The average (across-participants mean of the median per cost level) patch value differences (alternative - current patch) at which participants left their current patch increased with cost level (RM-ANOVA: F_3,81_ = 5.941, p = 0.001, *η*^2^ = 0.063, significant positive linear trend: t_27_ = 3.961, p < 0.001, CI_95_ = [1.341 – 4.223], Cohen’s U_3_ for one sample = 0.179; Figure 2A). To quantify how participants balanced patch values against cost, we computed a patch leaving advantage by subtracting, for every patch-leaving trial, the switch costs from the relative benefit of leaving (alternative - current patch value). These patch leaving advantages were then averaged across switch trials. The average patch leaving advantage across subjects was 19.39 ± 2.07 (mean ± SEM, see Figure 2B for an evolution of patch leaving advantages across all trials for one example participant).

**Figure 2.**
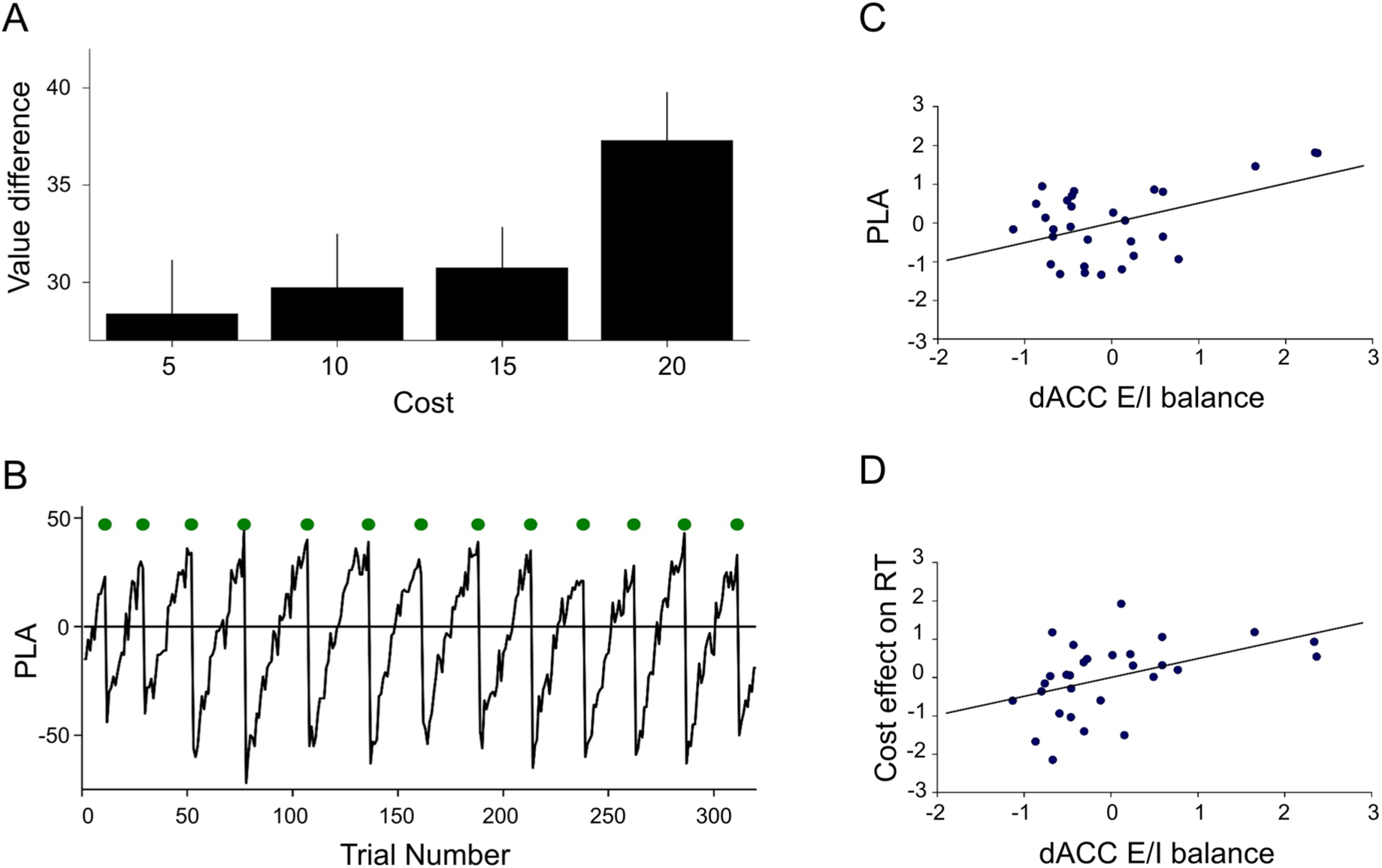
Patch-leaving behaviour and cortical E/I balance. A) Participants left their current patch at higher value differences (alternative - current patch value) when switching was associated with higher costs. Error bars indicate standard error of the mean. N=28. B) Example timecourse of patch leaving advantages (PLA) for one example participant. PLA = [value alternative patch] – [value alternative patch] – cost. Green circles indicate switch trials. C) Participants with higher dACC E/I balance leave at higher average PLA. N=29. D) Participants’ patch-leaving decisions are slowed down with increasing cost levels, and this effect is most pronounced in participants with high levels of dACC E/I balance. N=29.

To investigate the factors governing the speed of responding, we set up a multiple linear regression model. Patch value difference, patch-leaving trials, cost levels, trial number, switch (left/right) of patch presentation (relative to the previous trial), and wins in the previous trials were entered as independent variables to predict (log) response times. Participants’ responses were slower when switching entailed greater costs (t_28_ = 2.328, p = 0.027, CI_95_ = [0.001 – 0.016], U_3_ = 0.345), in switch trials (t_28_ = 3.397, p = 0.002, CI_95_ = [0.142 – 0.575], U_3_ = 0.276), when subjects had received reward at the value-guided choice stage of the previous trial (t_28_ = 3.249, p = 0.003, CI_95_ = [0.055 – 0.245], U_3_ = 0.207) and when there was a change in presentation sides of patch values (t_28_ = 2.960, p = 0.006, CI_95_ = [0.045 – 0.248], U_3_ = 0.276). Further to this, participants’ responding became significantly faster over the course of the experiment (t_28_ = −7.474, p < 0.001, CI_95_ = [-0.005 – 0.003], U_3_ = 0.897). There was no significant effect of patch value difference (t_28_ = 0.149, p = 0.882, CI_95_ = [-0.059 – 0.068], U_3_ = 0.517) on reaction times in the patch leaving phase.

### Cortical E/I balance and patch-leaving behaviour

To investigate how cortical E/I balance relates to patch-leaving decisions, we projected E/I balance (ratio between glutamate and GABA) in all five cortical areas onto behavioural variables of interest. To limit the number of statistical comparisons, we first ran this general linear model with the ratio of glutamate to GABA in all five areas. Only if this yielded a significant effect for one brain area, we then further tested for the individual contributions of GABA and glutamate to this effect by computing partial correlations, regressing out the effects of all other factors than the one currently of interest (see methods). Furthermore, where the general linear model yielded an effect of E/I balance for one region, we also report the partial correlation score for E/I balance in this region, since this is also what we used for visualization of the effect (figures 2C, D and 3C, D, E). We have computed a patch-leaving advantage that indicates how participants balance the relative benefit expected from leaving against the cost. Regressing E/I balance against patch leaving advantage revealed a significant influence of E/I balance in dACC (t_23_ = 2.643, p = 0.015, CI_95_ = [0.111 – 0.908]; r = 0.483, p = 0.008, CI_95_ = [0.141 – 0.722], Figure 2C) but not in any other region of interest (all p > 0.199). This effect was, by trend, driven by GABA in dACC (r = −0.323, p = 0.087, CI_95_ = [-0.617 – −0.049]; Supplementary Figure 1A), but not glutamate in dACC (r = −0.152, p = 0.431, CI_95_ = [−0.491 – 0.227]). In addition to that, a direct contrast showed that the relationship between patch leaving advantage and E/I balance was stronger in dACC compared to vmPFC (t_23_ = 2.613, p = 0.016) and dlPFC E/I (t_23_ = 2.087, p = 0.048). This finding suggests that the manner in which the relative benefit of leaving is balanced against travel costs is uniquely related to E/I balance in dACC, but not in any of the other areas investigated.

We next assessed how cortical E/I balance was related to patch response speed, and to how key task parameters affected response speed. We did not find any significant effects on overall response speed, but a specific effect of E/I balance in dACC on the degree to which patch decision choices were slowed by costs. Participants showed slowing of patch choices with higher cost levels, and the magnitude of this effect was significantly related to E/I balance in dACC (t_23_ = 2.461, p = 0.022, CI_95_ = [0.079 – 0.906]; r = 0.457, p = 0.013, CI_95_ = [0.108 – 0.705]; Figure 2D). This relationship was driven by GABA (r = −0.358, p = 0.056, CI_95_ = [−0.641 – 0.010], Supplementary Figure 1B) indicating that subjects with higher GABA levels in dACC show less increases in reaction times in relation to greater cost levels. None of the other task parameters that had a significant effect on reaction times were related to E/I balance (p > 0.143). Thus far, our results are consistent with our hypothesis. E/I balance in dACC, but not in any of the other regions investigated, is related both to how participants balance expected benefits against travel costs and to how costs affect the speed at which patch decisions are made.

### Value-guided choice behaviour

Participants selected the objectively correct option (higher expected value) in 81.85 % ± 1.52 (mean ± SEM) of all trials. We set up a logistic regression model to investigate the factors that affected participants’ decisions (right vs. left option). Participants choices were strongly guided by the options’ reward probabilities (t_28_ = 32.11, p < 0.001, CI_95_ = [0.949 – 1.080], U_3_ = 0), reward magnitudes (t_28_ = 16.161, p < 0.001, CI_95_ = [0.607 – −0.783], U_3_ = 0), and expected values (t_28_ = 9.363, p < 0.001, CI_95_ = 0.212 – 0.330], U_3_ = 0.069, Figure 3A). Probabilities had a greater effect on choice than magnitudes (t_28_ = 4.741, p < 0.001, CI_95_ = [0.181 – 0.470], d = 1.514), and both probabilities and magnitudes had a greater effect than expected values (t_28_ = 14.566, p < 0.001, CI_95_ = [0.639 – 0.848], d = 4.474; t_28_ = 13.935, p < 0.001, CI_95_ = [0.362 – 0.487], d = 2.110). In addition, participants exhibited a significant bias to choose the right option, over and above the effects of value parameters (t_28_ = 4.408, p < 0.001, CI_95_= [0.075 – 0.204], U_3_ = 0.138). There was no significant influence of either patch value difference (t_28 =_ - 1.386, p = 0.177, CI_95_ = [−0.088 – 0.017], U_3_ = 0.552) or switch costs (t_28_ = 0.501, p = 0.620, CI_95_ = [ −0.002 – 0.004], U_3_ = 0.448) in the patch-leaving phase on value-guided choice. Further, there was no significant effect of participants choices in the value guided choice phase of the previous trial (t_28_ = −0.167, p = 0.874, CI_95_ = [ −0.029 – 0.025], U_3_ = 0.448), of whether the left option had been rewarded in the previous trial (t_28_ = 1.214, p = 0.235, CI_95_= [ −0.012 – 0.047],U_3_ = 0.379) and of whether the right option had been rewarded in the previous trial’s value guided choice phase (t_28_ = 0.318, p = 0.753, CI_95_ = [ −0.028 – 0.038], U_3_ = 0.483, Figure 3A). Finally, there was no significant influence of the current trial’s patch leaving choice on the value guided choice (t_28_ = -1.654, p = 0.109, CI_95_ = [ −0.062 – 0.007], U_3_ = 0.621). This indicates that participants’ value-guided choices were indeed independent of the patch-leaving phase, and that choices were guided by the key value-related parameters, not by other aspects, such as whether or not a choice had been rewarded on the previous trial.

**Figure 3.**
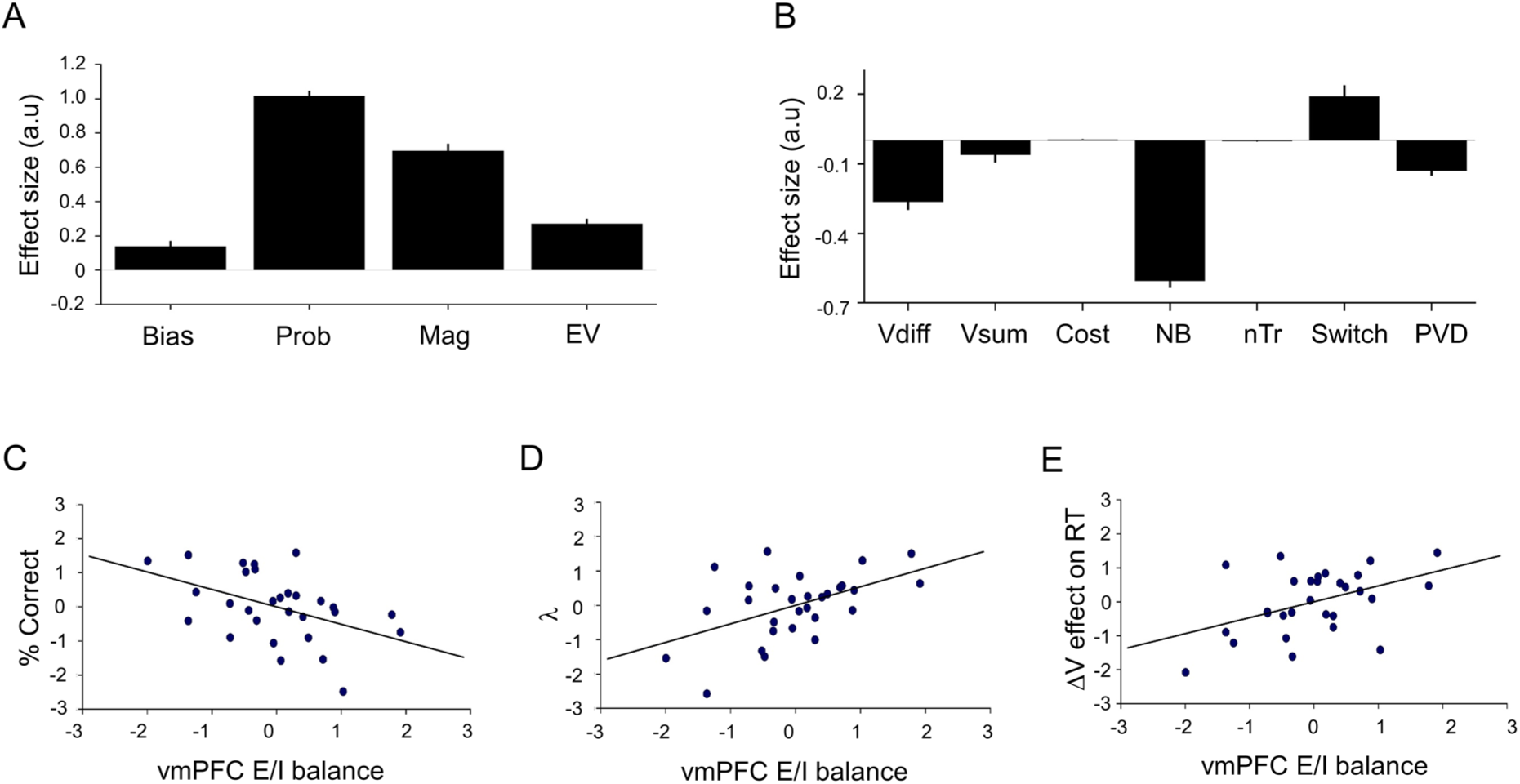
Value-guided choice behaviour and cortical E/I balance. A) Logistic Regression showing that participant’ choices (right vs left option) are guided by reward probabilities (Prob), magnitudes (Mag) and expected values (EV). B) Response times are significantly influenced by the value difference (Vdiff) between options, value sum (Vsum), no brainer trials (NB), trial number (nTr), patch value difference (PVD) and whether a trial is a patch-leaving trial or not (Switch). Error bars in A and B indicate standard error of the mean. C) Participants with high levels of E/I balance in vmPFC exhibit less accurate choice behaviour. D) Participants with high levels of E/I balance in vmPFC rely more strongly on differences in reward probabilities compared to reward magnitudes. E) Participants with high levels of E/I balance in vmPFC exhibit less slowing in difficult trials during value-guided choice. N=29.

To accumulate maximal returns, participants need to compute the Pascalian expected values by multiplying reward probabilities and reward magnitudes, and then choose the option with the higher expected value. However, humans do not weight probabilities and magnitudes in a statistically optimal way and show systematic distortions^8^. We fitted several different models, amongst them a standard prospect theory model^21^ (see methods), but we found that choices were best explained by a model in which values were computed as weighted combination of (weighted) attribute (probability and magnitude) differences and expected value differences:

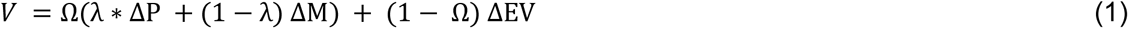

where ΔP, ΔM, and ΔEV are the differences in probability, magnitude and expected value, respectively. Ω and λ are two weighting parameters, both scaled between [0, 1]. Ω indicates the extent to which choices are guided by attribute comparisons as opposed to integrated Pascalian values, with high values of Ω indicating complete reliance on attribute comparison, whereas Ω = 0 would indicate perfect reliance on Pascalian values. Finally, λ indicates the degree to which attribute comparison is dominated by either probabilities or magnitudes. In essence, this model captures the findings from the logistic regression that showed that participants’ choices were guided by both attribute difference and expected value. Mean values across participants were Ω = 0.50 ± 0.05 and λ = 0.65 ± 0.04.

Finally, we investigated which factors affected the speed of value-guided decisions. We used multiple linear regression to test the effects of value difference, value sum and no-brainer trials (trials in which both probability and magnitude favoured choice of one option) on (log) response times. We additionally entered trial count, current patch value difference (alternative - current patch), patch leaving trials (1/0) and patch-leaving cost as control variables. Participants exhibited faster responding with greater value difference between chosen and unchosen option (t_28_ = -7.386, p < 0.001, CI_95_ = [−0.336 – −0.190], U_3_ = 0.966). Additionally, they exhibited a trend to faster responding in trials with higher value sum (t_28_ = -1.850, p = 0.075, CI_95_ = [ −0.129 – −0.007], U3 = 0.586). Furthermore, participants showed faster responding on trials with high patch value difference (t_28_ = -6.014, p < 0.001, CI_95_ = [ −0.174 – −0.086], U3 = 0.966), in no brainer trials (t_28_ = -20.790, p < 0.001, CI_95_ = [ −0.664 – −0.545], U3 = 1) and with increasing trial number (t_28_ = -7.830, p < 0.001, CI_95_ = [ −0.003 – −0.002], U3= 0.931). Finally, we found significantly slower responding in patch leaving trials (t_28_ = 3.842, p = 0.001, CI_95_ = [0.088 – 0.290], U3 = 0.207, Figure 3B). There was no effect of cost level (t_28_ = 1.370, p = 0.182, CI_95_ = [ −0.002 – 0.008], U_3_ = 0.379) on reaction times in the value-guided choice phase.

### Cortical E/I balance and value-guided choice behaviour

To relate cortical neurochemistry to value-guided choice behaviour, we used the same approach as above for the patch-leaving phase. Replicating our previous work^9^, we found that E/I balance in vmPFC was related to value-guided choice performance. Decision accuracy (% choices of the higher-value option) was negatively related to E/I balance in vmPFC (t_22_ = - 2.437, p = 0.023, CI_95_ = [−0.947 – 0.076]; r = −0.461, p = 0.012, CI_95_= [ −0.708 – −0.114], Figure 3C), but not in any of the other regions investigated (p > 0.406), indicating that subjects with higher concentrations of GABA relative to glutamate were better at selecting the higher value option. When we followed this up with partial correlations, neither GABA (r = 0.234, p = 0.221, CI_95_ = [ −0.145 – 0.553]) nor glutamate alone (r = −0.225, p = 0.241, CI_95_= [ −0.546 – 0.155]) was significantly correlated with decision accuracy. These findings indicate that participants with higher concentrations of glutamate in vmPFC indeed tend to exhibit less accuracy in their choice behaviour. We further tested whether E/I balance was related to how participants’ choices were guided by reward magnitudes and probabilities. We find a marginally significant effect of vmPFC E/I balance on the model parameter Ω (t_22_ = 1.923, p = 0.068, CI_95_ = [ −0.031 – 0.830]; r = 0.379, p = 0.042, CI_95_ = [0.015 – 0.655]), indicating that participants with higher concentrations of glutamate relative to GABA based their decisions more on attribute comparison than on integrated value. This effect was driven by GABA (r = −0.364, p = 0.052, CI_95_ = [ −0.645 – −0.003]). In addition to that, participants with high vmPFC E/I ratio also based their attribute comparisons more on probabilities compared to magnitudes, as indicated by a significant relationship between vmPFC E/I balance and λ (t_22_ = 2.726, p = 0.012, CI_95_ = [0.129 – 0.951]; r = 0.503, p = 0.006, CI_95_ = [0.167 – 0.734], Figure 3D). For both parameters, there was no effect of E/I balance in any of the other cortical regions (p > 0.114).The same pattern of results was obtained when we used the weights for probabilities and magnitudes, obtained from the logistic regression, as dependent variables. Again, we found participants with high vmPFC E/I ratio to be stronger influenced by probability differences than magnitude differences in their value-guided choice (magnitude weights: t_22_ = -2.832, p = 0.010, CI_95_ = [0.986 – −0.152]; r = −0.517, p = 0.005, CI_95_ = [ −0.743 – −0.186], Supplementary Figure 2A; probability weights: t_22_ = 2.120, p = 0.046, CI_95_ = [0.097 – 0.880]; r = 0.412, p = 0.026, CI_95_ = [0.053 – 0.676], Supplementary Figure 2B).

Taken together, value-guided choice performance was related to E/I balance in vmPFC, but not in any of the other cortical regions. Participants with high levels of GABA relative to glutamate were most reliable at selecting the higher-value options, based their choices more on integrated value than attribute comparison, and the extent to which they used attribute comparison was less skewed towards probabilities. Finally, we asked how value-guided response speed was related to cortical E/I balance. We first observed that log response times in the value-guided choice phase were specifically related to dACC E/I balance (t_22_ = - 2.337, p = 0.029, CI_95_ = [−0.965 – −0.058; r = −0.446, p = 0.015, CI_95_ = [ −0.698 – −0.095]). This effect was contributed to by a positive effect of GABA (r = 0.457, p = 0.013, CI_95_ = [0.108 – 0.705]), with no significant effect of glutamate (r = 0.026, p = 0.894, CI_95_ = [ −0.344 – 0.389]).

In contrast to these effects of dACC E/I balance on general response speed during value-guide choice, we found a specific effect of vmPFC E/I balance on the degree to which responses were speeded up by high value difference. The effect of value difference on response times in the value-guided choice phase was lowest in individuals with high vmPFC E/I balance (t_22_ = 2.388, p = 0.026, CI_95_ = [0.062 – 0.876]; r = 0.454, p= 0.013, CI_95_ = [0.105 – 0.703], Figure 3E). Specifically, GABA levels correlated negatively with this effect (r= −0.446, p= 0.015, CI_95_= [−0.698 – −0.095], Supplementary Figure 2C). This indicates that participants’ responses slowed down on difficult trials with low value difference, and that this slowing was most pronounced in individuals with relatively higher levels of GABA compared to glutamate. This pattern is consistent with our previous findings showing that vmPFC decision signals emerged more rapidly with higher concentrations of glutamate and low levels of GABA^9^. In addition, we found that the degree to which value-guided choices directly following a patch-leaving decision slowed down was by trend significantly related to E/I balance in left M1 (t_22_ = 1.922, p = 0.069, CI_95_ = [ −0.003 – 0.783]; r = 0.379, p = 0.043, CI_95_ = [0.015 – 0.655]). Specifically, increased GABA levels in left M1 were related to less slowing (r = −0.404, p = 0.030, CI_95_ = [ −0.671 – −0.044]) whereas the reverse effect was shown for glutamate levels in left M1 (r = 0.460, p = 0.012, CI_95_ = [0.113 – 0.707]).

In conclusion, this pattern of results mirrors the findings from the patch-leaving phase. Whereas patch-leaving behaviour was specifically related to dACC E/I balance, various parameters of value-guided choice behaviour were related to vmPFC E/I balance in a consistent and mechanistically plausible manner.

## Discussion

Knowing when to leave a depleting resource is a central problem for decision makers in naturalistic environments. It requires the agent to track the value of current resources, compare it to potential alternatives, and balance the potential benefits of moving against the cost incurred by moving. Within a given environment, it is crucial to consider the various attributes that jointly determine an option’s value - and to then select the most valuable option in order to maximize rewards. Thus, both patch leaving and value-guided decisions are key elements of adaptive behaviour. Using a novel behavioural task and assessment of cortical E/I balance by MRS quantification of GABA and glutamate concentrations, we have provided evidence for a double dissociation: E/I balance in dACC, but not any of the other regions investigated, was related to the manner in which participants balanced potential benefits of leaving against costs during patch-leaving decisions. In contrast, E/I balance in vmPFC was related to various aspects of value-guided choice.

Participants took costs into account in guiding their patch-leaving choices, as evident from the finding that they waited for higher advantages (higher value difference between current and alternative patch) as cost levels increased, and participants that required higher advantages compared to travel costs were characterized by high dACC E/I balance. Similarly, these participants also showed stronger slowing of patch response times with increasing cost levels. An extensive literature has implicated neural activity in dACC in behavioural adjustments^22–26^. Recently, these patterns of activity have been recast in light of new evidence suggesting that dACC may encode the evidence in favour of switching away from a current default option^11^. Specifically, dACC activity contained information about the value of searching the environment for better alternatives compared to the currently available options^3^. In both primates and rodents, firing of neurons in ACC ramps up just before the animal is about to abandon its current patch and move elsewhere^2, 4^, or when rats abandoned current beliefs and explored alternative strategies^27^. Similarly, ACC local field potentials in the gamma range have been related to switching between exploratory and exploitative modes of behaviour^27–29^. Since gamma oscillations are driven by a balance between glutamatergic excitation and inhibition by GABAergic interneurons^30–32^, it was plausible for us to assume a role for cortical E/I balance in patch-leaving decisions. We found that participants with higher E/I balance (higher levels of glutamate relative to GABA) required a higher patch leaving advantage (a higher difference between the benefits and costs expected from leaving) and showed more pronounced slowing of patch response times when costs were high. This effect showed a trend towards being driven by GABA, but not glutamate levels. Our findings are in line with previous reports showing a relationship between inhibitory neurotransmission in the dACC and patch-leaving behaviour. Parvalbumin-positive GABA interneurons in rodent anterior cingulate cortex have been shown to ramp up their firing prior to the animal leaving its current patch, and firing rates of these neurons represented the animal’s stay duration in the patch^4^. Furthermore, interhemispheric gamma synchronization driven by the same class of GABAergic interneurons in medial prefrontal cortex has recently been shown to enable mice to adaptively respond to changing environments^33^. Together with our results, these findings suggest that GABAergic activity in dACC may provide a signal for leaving one’s current patch. Another study in primates found a similar ramping pattern in dACC neurons^2^. While the cell type from which these recordings were obtained is not known, it is likely that they were predominantly obtained from glutamatergic pyramidal cells^34, 35^. This may appear contradictory at first glance, but could be easily reconciled when assuming that there is an asymmetry in the proportion of neuronal pools whose activity represents the value of leaving the current patch versus those that represent the value of staying. Such an assumption is plausible given that dACC has been shown to dominantly represent value of switching away from a current strategy^3, 7, 36^. Under such a scenario, in a recurrently connected network with GABAergic feedback inhibition, a ramping of pyramidal cell firing would recruit feedback inhibition, which would then further increase the asymmetry between the neuronal pools, gradually favouring the pools representing the value of switching. Thus, increased levels of GABAergic feedback inhibition would amplify the network transition towards favouring the alternative option and consequently bias the agent towards leaving the patch earlier. However, unlike for value-guided choice (see below), while a hypothetical model has been postulated^18^, to date there exist no biophysically plausible mechanistic models for patch-leaving behaviour.

In contrast to patch-leaving decisions, value-guided choice was specifically related to E/I balance in vmPFC: high vmPFC concentrations of GABA relative to glutamate were related to (1) an increased decisions accuracy (selection of the higher-value option), (2) a stronger reliance on integrated value as compared to attribute comparison, and (3) the extent to which participants relied on attribute comparison was less biased towards reward probabilities. Furthermore, vmPFC GABA concentrations were also related to how participants slowed on difficult trials (choices with low value difference), with participants with high GABA concentrations again showing more pronounced slowing. These results are in line both with mechanistic models of decision making^17, 37^ and our own previous findings^9^. It is thought that decisions may be generated by a mechanism that is based on competition via mutual inhibition in recurrent cortical networks that exhibit attractor dynamics^37, 38^. In these models, a winner-take all competition is implemented, where (in the binary case), activity in only one of the two pools representing the two options remains (the chosen option), whereas activity in the other pool is suppressed. One key prediction of these models is that increased GABAergic feedback inhibition slows down the attractor dynamics, allowing for more evidence to be accumulated^9, 17^. Thus, increased GABAergic tone makes decisions slower but more accurate. In our previous work, we showed that higher concentrations of GABA and low concentrations of glutamate were related to increased decision accuracy. Neurally, this was accompanied by a slower, but more stable ramping of a value difference correlate in vmPFC, a neural signature of a decision^9^. Our present results exactly match with this pattern. Choice performance was highest in participants with high vmPFC concentrations of GABA relative to glutamate, and these participants also showed the most pronounced slowing on difficult trials. As reported above, task-specific effects were found for patch-leaving and value-guided choice in dACC and vmPFC, respectively. However, while participants were significantly influenced by value differences between choice options during value-guided choice, we did not find any significant influence of patch value difference on reaction times during patch-leaving decisions. This discrepancy between the two stages may appear surprising at first glance. However, response times likely indicate rather different factors in the two stages. In the patch stage, participants can already make up their mind whether to switch or stay on the next trial immediately after they observe the outcome of their patch choice. In contrast, at the value-guided choice, participants cannot anticipate the options they will encounter and instead have to compute option values “on the fly”.

A notable aspect of our findings is that response times in each phase were modulated by events from the respective other phase. Participants’ value-guided choices speeded up as the value difference between the alternative and current patch increased on trials leading up to a switch, but when participants chose to leave their patch, the immediately subsequent value-guided choice was slowed down. Conversely, patch-leaving decisions were slowed when the previous trial’s value-guided choice had been rewarded. The functional significance of these effects is not clear, but the former might indicate that participants switch to a more cautious, evaluative mode of value-guided choice upon entering the alternative patch. A recent modeling account^36^ suggests an interplay between dorsomedial prefrontal cortex (including dACC) and vmPFC in deciding when to switch away from ongoing behaviour, based upon reliability ratings of the current strategy^39^. In our task, rewards are only obtained during value-guided choice. These potentially serve as a feedback on the current strategy, which in turn might mediate a switch from ongoing behaviour. It has been suggested previously^40^ that increased glutamate levels in dACC lead participants to exploit underlying task structure, whereas increased GABA concentrations allow for learning a new model of that task.

In summary, we have shown that cortical E/I balance, as assessed by MR spectroscopic quantification of baseline GABA and glutamate concentrations is related to both patch-leaving and value-guided decision making. We found a double dissociation, where E/I balance in dACC is related to patch-leaving, but E/I balance in vmPFC is related to value-guided choice. The pattern of results further supports models that implement a competition via mutual inhibition in recurrent cortical networks as a candidate mechanism for value-guided choice. Importantly, we provide novel evidence that relates dACC E/I balance to patch-leaving decisions. The pattern of results suggests that elevated GABAergic relative to glutamatergic tone in dACC may increase the propensity to switch away from a current policy. Understanding the neurochemical mechanisms underlying different types of decision making is of potential clinical relevance, since alterations in E/I balance have been described in a number of neuropsychiatric disorders^41, 42^ which are characterized by impaired decision making behaviour^43, 44^.

## Methods

### Participants

Thirty-three right-handed (Oldfield-Score^45^: 91.95 ± 1.89, mean ± SEM) male participants (age: 26.18 ± 0.65, mean ± SEM, range: 22-36) with normal (N = 16) or corrected to normal (N = 17) vision participated in this experiment. Exclusion criteria comprised a history of neurological or psychiatric illness, drug abuse and use of psychoactive drugs or medication 24 h prior to participation. Four subjects were excluded due to excessive noise in at least one of five spectroscopy measurements (see MR imaging for criteria). All reported results are from the remaining N = 29 subjects (mean age: 26.48 ± 0.72, range: 22-36; normal vision: N = 14; non-smoker: N = 22). Written informed consent to the procedure was obtained from all subjects prior to the experiment, which was approved by the local ethics committee of the Medical Faculty of the Otto-von-Guericke-University, Magdeburg. Participants were compensated for each session and received a bonus that depended on their performance in the decision making task.

### General study procedure

Each participant was measured twice (average time between sessions: 1.52 days). In the first session, participants were asked to participate in a decision making task. During this task, participants were scanned with Magnetoencephalography (MEG, data not presented here). In the second session, participants underwent scanning with MR spectroscopy at 7T for quantification of cortical GABA and glutamate concentrations^46^.

### Decision making task

Participants were asked to maximize their rewards in a two stage decision making task consisting of 320 trials (Figure 1A). They first completed 15 practice trials to familiarize themselves with the task before commencing the experiment. Stimulus presentation was controlled by Psychtoolbox 3^47, 48^ running on Matlab 2012b (The Mathworks Company, Natick, MA). Each trial started with presentation of the two patches (two grey squares). The patch in which the participant currently resided was indicated by a grey frame around the patch. At this stage, participants simply had to indicate by button press (with the index finger of the left or right hand, respectively) whether they wanted to stay in their current patch or switch to the alternative patch. If participants chose to leave their patch, they had to pay a travel cost indicated by the size of a grey bar presented centrally between the two patches. Travel costs were randomly drawn from the set {5, 10, 15, 20 points} and remained constant until a participant chose to leave their patch, at which stage a new cost was selected. The participant’s patch choice was highlighted by a frame around the selected patch (400 - 600 ms, jittered). In trials where participants chose to leave, the rectangular bar signaling travel costs turned red and the costs were subtracted from a progress bar displayed below the patches that indicated the participant’s total earnings. Presentation sides (left/right) of the two patches were randomly selected on each trial. Therefore, while participants could decide in advance whether they wanted to stay or leave their patch, this prevented them from preparing the actual motor response before trial onset. Afterwards, the values of the two patches (the reward available in each of them, as indicated by the blue filling) was revealed (1800 - 2200 ms, jittered). Importantly, the value of the participant’s current patch stochastically depleted over time, whereas the alternative patch replenished. Therefore, participants were required to continually accumulate evidence in favour of abandoning their current patch. On each trial, values of the two patches were drawn from Gaussian distributions with non-stationary means and variance = 3.5. The means μ of both patches were set to 50 points initially and then diffused according to a decaying Gaussian random walk on each trial:

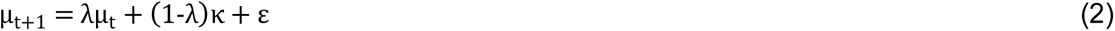

where λ is the decay rate which was set to 0.96, κ is the decay centre (1 for the chosen patch, 100 for the unchosen patch) and ε is zero-mean Gaussian random noise with a standard deviation = 1.2. If a patch value drawn from these distributions was above an upper bound of 90 or below a lower bound of 10 points, the value was set to this bound. After the value of the two patches was revealed, participants entered the second stage value-guided choice. The reward available in the chosen patch was allocated to two choice options at a random ratio (ensuring that none of the two options received less than 10% of the total patch value and excluding a 50% split between options). Furthermore, both options were assigned a probability with which this reward could be obtained, randomly sampled from the set {0.1, 0.2, …, 0.9}. Reward probabilities were independent of each other, such that in any given trial, either of the two options, both options, or neither of them could be rewarded. Importantly, while higher patch values will, on average, lead to better choice options, this procedure ensures that participants do not know the two choice options when they make their patch decision, thereby explicitly decoupling the patch choice from the value-guided choice. In trials where both the reward probability and magnitude of one option was higher than that of the other option (a ‘no-brainer trial’), we randomly flipped the reward magnitudes of the two options in 50% of cases to control for task difficulty. Due to an error in our code however, this change was only applied to no-brainer trials in which the left option had higher values than the right one. Reward magnitudes were indicated by the height of a grey bar and reward probabilities were presented as numbers (in percent) below each bar (Figure 1A). Reward magnitudes were displayed relative to an outline that corresponded to the overall reward available in the current patch. Participants selected an option by pressing a button with the right or left index finger, respectively. The chosen option was highlighted by a rectangular frame and the outcome of both options was revealed (800 – 1200 ms, jittered). The bar representing the reward magnitude turned green if an option was rewarded in the current trial, or red otherwise. Even though participants could not benefit from knowing whether the unchosen option would have been rewarded, this procedure has proven useful to remind participants that even low probability options are occasionally rewarded and that it is beneficial to consider each option’s reward probability and magnitude. Every time participants were rewarded, the progress bar grew (in proportion to the obtained magnitude) towards a goal indicated by a gold target line to the right of the screen. Outcome presentation was followed by an intertrial interval (1800 - 2200 ms, jittered) before participants entered the patch decision stage of the next trial. A centrally located fixation cross was present throughout the entire trial. See supplementary materials for further details on stimulus presentation.

### Analysis of behavioural parameters

For the patch-leaving phase, we computed, for each participant and each cost level separately, the median patch value difference (alternative – current patch) at which participants left their current patch. These average patch value differences were compared across cost levels using a RM-ANOVA. Additionally, linear trends along with a constant term were regressed against cost-level dependent switching behaviour to estimate whether the median patch value difference increased linearly with cost level. In all cost-level dependent analyses, the data of N = 28 participants were analyzed since one subject was never presented with the highest cost level (cost levels were randomly assigned after each switch). Furthermore, we defined a patch-leaving advantage by subtracting travel costs from patch value differences at each patch-leaving decision. These values were then averaged across switch trials per participant. We used patch value differences from the previous trial for all analyses pertaining to the patch-leaving phase since the updated patch values are only revealed following the patch choice and hence are informative for the next trial.

For the value-guided choice phase, we computed the percentage of correct responses as the percentage of trials in which subjects chose the option with higher expected value.

Regression coefficients were tested against zero with a t-test for one sample (two-sided).The MEST toolbox was used to provide estimates for effect size measures (η^2^ for repeated measures ANOVA and Cohen’s U3 for one sample to test regression coefficients against 0)^49^. Additionally, 95 % confidence intervals for the mean of regression coefficients across subjects are reported.

### Regression analyses

For each regression analysis, all continuous variables were normalized (z-scored) and a constant term was added to each design matrix. All regressions were performed for each participant separately, and regression coefficients were normalized by their vector norm before averaging across subjects. Regression coefficients were tested against zero using one-sample t-tests (two-tailed) and the MEST toolbox was used to provide estimates for effect size measures (Cohen’s U3 for one sample, Cohen’s d for paired samples)^49, 50^. Additionally, we report the 95% confidence interval for the mean of each distribution of regression coefficients (or their differences in the case of paired samples) across participants.

To analyze how key value parameters influence value-guided choice, we set up a multiple logistic regression model with choice of the left vs right (0/1) option as dependent variable. Probabilities and magnitudes for both options, expected values (the product of demeaned probabilities and demeaned magnitudes for each option) as well as patch value differences and travel costs were entered as independent variables. Here, we used patch value differences from the current trial since they are already known by the participant at the time of their value-guided choice. Furthermore, we added the previous trial’s value guided choice, the current trial’s patch leaving choice, and two additional regressor coding for whether the right and left option, respectively, had been rewarded in the previous trial to the design matrix. A paired sample two-sided t-test was used compare between regression weights. To test the effects of value-guided choice parameters (probability, magnitude and expected value) against zero, regression weights for the left option were subtracted from those for the right option.

To analyze how various task parameters influenced response speed both in the patch-leaving and value-guided choice stage, we used multiple linear regressions with log response time as the dependent variable. As above for the logistic regression, continuous variables were normalized, a constant was added to the design matrix, regressions were run for each participant separately, and coefficients were normalized by the within-subject vector norm before averaging across participants. For the patch stage, the design matrix included patch value differences from the previous trial, a binary regressor indicating whether the current trial was a switch trial (one in which the participant left their current patch), the travel cost, a regressor coding for the linear effect of trial number, and two binary regressors indicating whether the side (left/right) of patch presentation had changed with respected to the previous trial and whether the previous trial’s value-guided choice had been rewarded. We used the patch value difference from the previous trial, because the outcome of the (updated) patch values are only revealed after participants’ patch choice. For the value-guided choice stage, the design matrix included the value difference between the chosen and unchosen option of each trial, the value sum of both options, travel costs, a binary regressor indicating ’no-brainer’ trials, a regressor coding for the linear effect of trial number, a binary regressor indicating whether the current trial was a patch-leaving trial and one regressor containing the patch value difference from the current trial.

### Behavioural modelling of value-guided decisions

Since it is known that humans do not weight magnitudes and probabilities in a statistically optimal way, we derived subjective magnitudes and probabilities by fitting utility functions according to prospect theory^51^ to each subject’s choice data in the value-guided choice phase. Each trial’s objective reward magnitudes and probabilities were transformed to subjective reward magnitudes and probabilities according to the following equations:

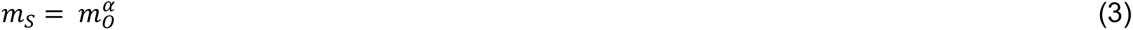

where m_S_ is the subjective reward magnitude, m_O_ is the objective reward magnitude and α is a free parameter used to fit the subjective magnitude. Subjective probabilities were estimated by:

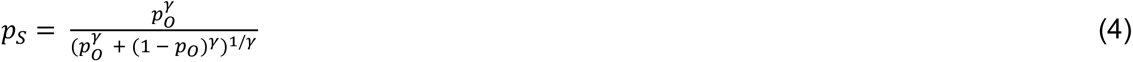

where p_s_ are the subjective reward probabilities, p_O_ are the objective reward probabilities and γ is a free parameter used to fit subjective reward probabilities. These values can be combined into subjective expected values by multiplying subjective reward probabilities and subjective reward magnitudes. Based on the expected values, choice probabilities were modeled with a modified softmax choice rule^52^

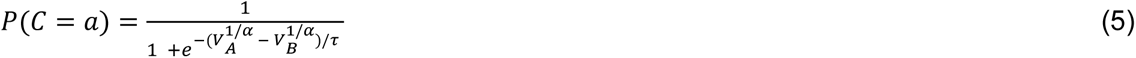

Where τ is a temperature parameter that describes the extent of stochasticity in participants’ choice behaviour. V_A_ and V_B_ are the expected values for the two options (the products of subjective magnitudes and subjective probabilities). The search space for each free parameter was constrained between 0 and 3. In addition to this full three parameter prospect theory model, we also fitted reduced parameter models that either lacked magnitude or probability distortion (α or γ set to 1, respectively) or lacked both magnitude and probability distortion (α and γ set to 1) to test whether these more parsimonious models provide a better account of participants’ choice data. The Bayesian Information Criterion (BIC) was used to compare between different models (see supplementary table S1 for an overview of fitted model parameters).

An alternative manner in which participants might construct option values is by means of a weighted attribute comparison:

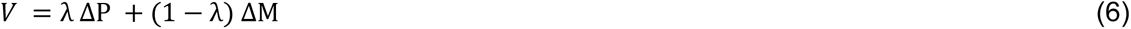

where ΔP and ΔM are the differences in probability and magnitude, respectively. The weighting parameter λ indicates the degree to which attribute comparison is dominated by either probabilities or magnitudes. Finally, it is also possible that values result both from a weighted sum of the above (weighted) attribute comparison and integrated (Pascalian) value:

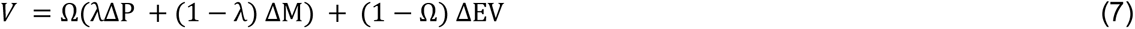

where ΔEV is the differences in expected value and Ω is a weighting parameter that indicates the extent to which choices are guided by (weighted) attribute comparisons as opposed to integrated Pascalian values. The space for both weighting parameters, λ and Ω, was constrained between 0 and 1. For both the attribute comparison only model and the attribute comparison + integrated value model, choice probabilities were modeled with a standard softmax choice rule:

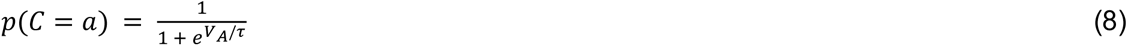

Probabilities, magnitudes and expected values were rescaled between 0.1 and 1 before model fitting. Parameters were optimized using custom-written scripts in MATLAB R2019a (The Mathworks Company, Natick, MA) and constrained non-linear optimization using MATLAB’s function fmincon to minimize the negative log likelihood of the data given the parameters. In order to decrease the probability of fitting local minima, we used 10,000 random starting points and report the combination of parameters with the lowest negative log likelihood.

### MRS data acquisition

MR data were acquired on a 7T system (Siemens Healthineers) equipped with a 32 channel array head coil (Nova Medical). First, a high-resolution T1 weighted scan was acquired using an MPRAGE sequence (TE = 2.73 ms, TR = 2300 ms, TI = 1050 ms, flip angle = 5°, bandwidth = 150 Hz/pixel, acquisition matrix = 320 x 320 x 224, voxel size = 0.8 mm³ isotropic) aligned with the anterior-posterior commissure. This scan was used for placement of MRS voxels, but also for tissue segmentation. We positioned voxels in five regions of interest, including right dorsolateral prefrontal cortex (dlPFC), bilateral primary motor cortices (rM1 and lM1), perigenual anterior cingulate cortex within vmPFC (vmPFC/pgACC) and dorsal anterior cingulate cortex (dACC). The dlPFC voxel was placed on the right hemisphere within the middle frontal gyrus by using the superior frontal sulcus and the inferior frontal sulcus as anatomical landmarks. We positioned the voxel as far dorsally as possible when excluding the calvaria and all extracalvarcial structures. The average dlPFC voxel centroid across participants was estimated at MNI x = 29.79 ± 0.85, y = 37.72 ± 1.38, z = 24.21 ± 1.51 (mean ± SEM). Primary motor cortex voxels were placed on the hand knob structures, identified by their omega-like shape on the central sulcus in axial slices. Average M1 voxel centroids in standard space were estimated at MNI x = -28.97 ± 0.82, y = -18.48 ± 0.92, z = 51.86 ± 0.59 and MNI x = 31.90 ± 0.71, y = -14.76 ± 1.06, z = 49.76 ± 0.88 for rM1 and lM1, respectively. The vmPFC voxel was mediolaterally centred on the midline and dorsoventrally on the genu of the corpus callosum, with its posterior boundary just rostral to the genu. The average voxel centroid position across subjects was estimated at MNI x = −0.17 ± 0.15, y = 41.41 ± 1.29, z = 7.00 ± 0.44. The dACC voxel was placed with reference to the corpus callosum, the cingulate as well as surrounding sulci. We used the posterior border of the genu of the corpus callosum perpendicular to AC-PC orientation to centre the voxel (Figure 1B)^46^. The average centroid voxel position across subjects was MNI x = −0.07 ± 0.19, y = 24.14 ± 0.43, z = 29.69 ± 0.47 (Figure 1B). For the MRS measurements, region specific shimming was performed using a double-gradient echo shim technique. Voxel sizes were 10 x 20 x 15 mm³ for the vmPFC voxel and 10 x 25 x 15 mm³ for all other voxels of interest. Afterwards, MR spectra were acquired using a stimulated echo acquisition mode (STEAM VERSE) sequence (128 averages, TR = 3000 ms, TE = 20 ms, mixing time = 10 ms, data size = 2048, bandwidth = 2800 Hz) from each voxel of interest^46^.

### MR Data Analysis

Spectral data were analyzed using LC Model^53^. Only metabolite measurements with a Cramér– Rao lower bound (CRlb) < 20 %, full-width half-maximum (FWHM) < 25 Hz and signal-to-noise ratio (SNRs) > 8 were included. We analyzed the quality of each voxel measurement using LCModel immediately after acquisition of the voxel. If one voxel did not meet the quality criteria, we repeated the acquisition of this specific voxel. We had to repeat measurement of one of the five voxels in 13 of our 29 subjects to obtain valid measurements for all five voxels of interest. SPM 12 (Wellcome Trust Centre for Neuroimaging, London, United Kingdom) was used to segment participants T1-weighted anatomical images into grey matter (GM), white matter (WM), cerebrospinal fluid, soft tissue and air/background. Each voxel’s GABA and glutamate concentrations were corrected for relative gray matter (GM) concentrations^54^ by dividing their absolute concentrations by relative GM, based on the assumption that GABA and glutamate are predominantly present in GM. As SPM 12 provides tissue probability maps, we summed across probabilities for grey matter for each voxel in the mask and divided by the total number of voxels within each mask to approximate relative GM. Total creatine concentrations (creatine + phosphocreatine) were normalized by the relative amount of gray matter and white matter within each voxel ((GM+WM)/number of voxelsmask) based on the assumption that creatine is predominantly present in GM and WM. All GABA and glutamate concentrations we report are normalized by total creatine concentrations. We defined E/I balance as the ratio of (normalized) glutamate to GABA levels. Voxel masks were then interpolated to individual MRI volumes with FieldTrip^55^. To estimate average voxel centroid positions we normalized individual volume data. More specifically, data were registered to MNI space by using tissue probability maps (TPM.nii template from SPM 12). Estimated average centroid positions were extracted from each mask in MNI space.

### Behavioural parameters and E/I balance

From each behavioural analysis we obtained estimates regarding the individual influence of key value parameters on behaviour (decision variables). To reduce the number of multiple comparisons when testing model parameters for its relationship with neurochemistry, we combined glutamate and GABA to one value by dividing glutamate by GABA concentrations, referred to as E/I balance. The ratio of glutamate to GABA within all voxels of interest were entered as linear regressors in a design matrix (along with a constant term) to predict the contributions of E/I balance onto each decision variable (one linear model per decision variable). In addition to t- and p-values we report the 95 % confidence intervals for each significant (p < 0.05) regression coefficient. Only if a significant influence of E/I balance was measured, partial correlations for E/I balances were estimated for the respective region. Therefore, E/I balances within the respective region as well as the decision variable were orthogonalized successively with respect to all other E/I balances (regression used for orthogonalization without constant term). The residuals of the decision variable and residual E/I balances within the respective region of interest where then correlated (Pearson correlation). We report the correlation results along with their respective p-values and 95 % confidence intervals for each correlation coefficient. Further, we estimated contributions of GABA and glutamate separately given that we found a significant effect of E/I balance for this region. Therefore, glutamate and GABA concentrations within the respective region (as well as the decision variable) were orthogonalized successively with respect to all other neurotransmitters (GABA or glutamate, respectively, within the same voxel, and GABA and glutamate in all other voxels). In the analysis of value-guided decisions, we also controlled for the amount of no brainer trials (mean ± SEM: 38.52 % ± 0.01) as a predictor variable of no interest. The residuals of the decision variable and residual GABA or glutamate concentration within the respective region of interest where then correlated (Pearson correlation). We report the respective r- and p-values for each correlation as well as the 95 % confidence interval for each regression coefficient.

## Data availability

The MRS data that support the findings of this study are available from the corresponding author upon reasonable request. The behavioural data are available under www.github.com/luckyluc25/clash_exp

## Code Availability

Custom written code used to analyze the behavioural data of the current study is available under www.github.com/luckyluc25/clash_exp

## Acknowledgments

The authors thank Renate Blobel-Lüer for her help with MRS recordings.

## Author Contributions

TOJG & GJ designed the research. TOJG & OS recorded the data. LFK, TOJG and GJ analyzed the data. LFK & GJ wrote the manuscript. LFK, TOJG, LL, OS and GJ read and edited versions and approved the final version of the manuscript. All authors discussed the results at all stages of the experiment.

## Competing interests

The authors declare no competing financial interests.

