## Supplementary Material for "Dissociable roles of cortical excitation-inhibition balance during patch-leaving versus value-guided decisions"

### Supplementary Materials

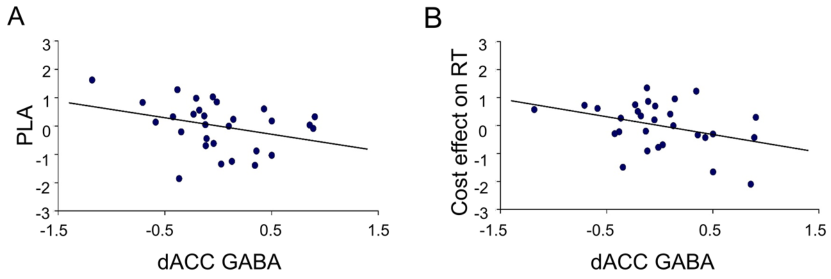


**Supplementary Figure S1:** GABA contribution to the effect of E/I balance on patch-leaving shown in main figure 2. A) Higher dACC concentrations of GABA are associated with earlier patch leaving (lower average patch leaving advantages. B) Participants' patch-leaving decisions are slowed down with increasing cost levels, and this effect is more pronounced in participants with low dACC GABA concentrations. N=29.

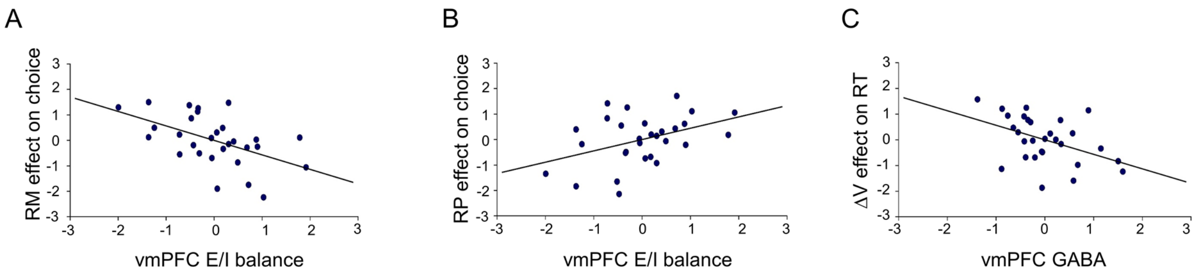


**Supplementary Figure S2:** Higher levels of vmPFC E/I balance were related to diminished effect of reward magnitudes (RM, A), but increased effect of reward probabilities (RP, B) on value-guided choice, indicating a bias towards choices being dominated by probabilities versus magnitudes. C) GABA contribution to the effect of E/I balance shown in main figure 3E. Participants' responses slowed down on difficult trials (trials with low value difference). This effect was related to vmPFC GABA concentrations. ΔV = value difference.

**Supplementary table 1: Parameter values and model fits for different behavioural models**

| Model | α | γ | τ | Ω | λ | BIC |
|---|---|---|---|---|---|---|
| α | 0.53 ± 0.06 | | | | | 199.58 ± 12.31 |
| α, τ | 0.58 ± 0.06 | | 1.71 ± 0.14 | | | 182.11 ± 9.46 |
| α , γ | 0.62 ± 0.10 | 1.13 ± 0.16 | | | | 182.75 + 9.65 |
| γ | | 1.99 ± 0.20 | | | | 239.53 ± 20.07 |
| γ,τ | | 2.13 ± 0.15 | 2.22 ± 0.14 | | | 192.23 ± 11.38 |
| τ | | | 2.72 ± 0.11 | | | 299.17 ± 30.27 |
| α,γ, τ | 0.59 ± 0.06 | 1.09 ± 0.12 | 1.26 ± 0.11 | | | 181.13 ± 10.10 |
| λ | | | | | 0.71 ± 0.08 | 379.92 ± 2.17 |
| Ω | | | | 0.00 ± 0.00 | | 399.29 ± 1.81 |
| λ , τ | | | 0.06 ± 0.00 | | 0.53 ± 0.02 | 184.41 ± 6.60 |
| Ω, τ | | | 0.07 ± 0.01 | 0.34 ± 0.00 | | 251.91 ± 14.22 |
| Ω, λ | | | | 0.89 ± 0.04 | 0.75 ± 0.08 | 385.30 ± 2.12 |
| Ω, λ ,τ | | | **0.05 ± 0.00** | **0.50 ± 0.05** | **0.65 ± 0.04** | **169.83 ± 7.12** |

*Note*: All values are mean values across participants ± standard error of the mean. N=29.

#### Supplementary Methods

##### Details of the behavioural task

All stimuli were presented on a grey (RGB: 60, 60, 60) background with a contrast optimized for the MEG recording chamber on a screen in a distance of one meter from the sitting participants. Stimuli were displayed via a projector with a refresh rate of 75 Hz located outside the MEG recording chamber. During patch-leaving, participants were presented with two patches (RGB: 80, 80, 80) framed with a white outline indicating in which patch participants are currently staying. If participants chose to switch they had to pay a travelling cost indicated by the size of a grey bar (RGB: 160, 160, 160) presented between both patches. In trials where participants chose to switch, the rectangular bar signaling switch costs turned red (RGB: 178, 70 70) and the respective costs were subtracted from the subjects total earnings up to this trial. Afterwards the current patch values were revealed. Patch Values were presented in blue (RGB: 69,102,174). In both stages of the experiment a blue progress bar (RGB: 65,105,204) was shown at the bottom of the screen indicating subjects current score. Participants selected an option by means of a button press with the right or left index finger, respectively. After value-guided choice, participants received a feedback on both options. If an option was rewarded in the current trial, the bar presenting the reward magnitude turned green (RGB: 46, 139, 60) or red (RGB: 178, 70, 70) otherwise. Every time participants were rewarded, the progress bar grew proportional to the obtained magnitude towards a goal state indicated by a golden rectangle (RGB: 184, 134, 11). The goal in the experiment was to reach the goal state as often as possible.
